# Assessment of high-efficacy agonism in synthetic cannabinoid receptor agonists containing l-*tert*-leucinate

**DOI:** 10.1101/2024.10.11.617959

**Authors:** Christopher Lucaj, Soo Jung Oh, Charlotte Pitha, Jordan Davis, Hideaki Yano

## Abstract

Synthetic cannabinoid receptor agonists (SCRAs) represent a class of new psychoactive substances that pose great health risks attributed to their wide-ranging and severe adverse effects. Recent evidence has shown that SCRAs with key moieties can confer superagonism, yet this phenomenon is still not well understood. Here we report structure-activity relationships (SARs) of modular SCRAs contributing to superagonism by comparing eight compounds differing by their head moiety (l-valinate vs. l-*tert*-leucinate), core moiety (indole vs. indazole), and tail moiety (5-fluoropentyl vs. 4-fluorobenzyl) through different modes of bioluminescence resonance energy transfer (BRET) assays. We found that the l-*tert*-leucinate head moiety and indazole core moiety conferred superagonism across multiple Gα_i/o_ proteins and β-arrestin-2. After generating the cannabinoid type 1 receptor (CB1R) mutant constructs, we found that transmembrane 2 (TM2) interactions to the head moiety of tested SCRAs at F170, F174, F177, and H178 are key to eliciting activity. Finally, we found that l-*tert*-leucinate SCRAs confer a high-efficacy response in *ex vivo* slice electrophysiology.

## 1. Introduction

The severe adverse effects observed with SCRA use are distinct from those of delta-9-tetrahydrocannabinol (THC), the active compound found in *Cannabis sativa* [1–4]. Due to the aminoalkyl-heterocycle scaffolds of SCRAs, drugs can be made through iterative modular design, resulting in greatly increased efficacy and potency and leading to their illicit use, as noted by the >100 novel SCRAs identified by the United Nations Office on Drugs and Crime every year since 2013 [5]. MMB-FUBINACA, one of these SCRAs, was responsible for a well-publicized case of SCRA intoxication that occurred in 2016 when thirty-three people exhibiting “zombie-like” cataleptic behavior [6]. In a separate clinical case, patients experienced extreme agitation, aggressiveness, and seizures after using another SCRA ADB-PINACA [3]. Compared to chronic cannabis use, cessation of SCRA use can lead to more severe withdrawal symptoms, with less frequency and volume of consumption [7, 8]. Although rare, SCRA overdose lethality has been reported, whereas that has not been reported with cannabis use [9–12].

With high affinity, SCRAs bind CB1R, a highly expressed G-protein coupled receptor (GPCR) in the brain and CNS [1, 13]. CB1R plays a crucial role in downregulating neurotransmitter release, primarily through Gα_i/o_ signaling pathways initiated by retrograde signaling at synaptic terminals [14, 15]. Upon G-protein activation, voltage-gated Ca^2+^ channels are inhibited and G-protein-gated inward rectifying K^+^ channels are activated, resulting in pre-synaptic hyperpolarization [14, 16]. Peripherally, the receptor also plays an important role in cardiovascular function as well as energy metabolism [17–19]. CB1R expression and function in multiple systems are complemented by the differential expression of the Gα_i/o_ protein subfamily, consisting of ubiquitously (Gα_i1_, Gα_i2_, Gα_i3_) and neuronally (Gα_oA_, Gα_oB_, Gα_z_) expressed inhibitory G-proteins [20–23]. Pharmacological studies have characterized a number of SCRAs based on CB1R-mediated cAMP inhibition as well as β-arrestin internalization [24–27]. SCRA-induced G-protein signaling within the subfamily of Gα_i/o_ proteins, however, has only been investigated very recently [26, 28, 29]. Though these studies found SCRAs to show little functional selectivity or bias, a thorough comparison of moieties among structural analogs has not been attempted by using rigorous proximity assay approaches.

With the increasing number of newly reported chemical structures, SCRA pharmacology has been assessed by potency and efficacy of conventional functional assays. Indeed, SARs of these chemical structures, composed of three key moieties (head, core, and tail), have led to a characterization of SCRAs as “high efficacy” agonists [6, 10, 24, 25]. However, assays that measure activity downstream of receptor-transducer interactions may be affected by signal amplification, leading to an inaccurate assessment of reduced efficacy and increased potency [30]. This limitation warrants the incorporation of a rigorous study design that is less affected by signal amplification. Recently, we investigated the SAR between two compounds, 5F-MMB-PICA (M-PC) and 5F-MDMB-PICA (D-PC), that differed in a single methyl group in the “head” moiety [31]. D-PC, with an l-*tert*-leucinate head moiety, was shown to act as a “superagonist”, an agonist that has greater efficacy than that of the endogenous ligand, while M-PC acted as a full agonist [30, 31]. Notably, molecular dynamics simulations between these drugs revealed different levels of interaction with key residues in the extracellular TM2 domain, a region recently reported to be critical in activation of CB1R [31–34]. Although we found a SCRA that elicits superagonism, an understanding of what drives CB1R superagonism is still limited.

In the current study, we further explore differences in SCRA moieties to uncover CB1R superagonism based on differences in “head,” “core,” and “tail” moieties. With a panel of eight compounds, we use different modes of BRET assays to assess the efficacy, bias, and functional selectivity of these compounds. We then introduced point mutations in TM2 residues of CB1R to reveal key interactions with the head moiety l-*tert*-leucinate. Finally, we measured the effects of SCRAs on CB1R-mediated inhibition of glutamate release in the hippocampus through *ex vivo* slice electrophysiology.

## 2. Materials and Methods

### 2.1 Mammalian Cell Culture

All *in vitro* assays are performed in human embryonic kidney 293 T (HEK-293T) cells cultured in Dulbecco’s Modified Eagle Medium (DMEM) supplemented with 10% fetal bovine serum (FBS), 1% penicillin/streptomycin, and 2 mM L-glutamine and incubated at 37°C and 5% CO_2_. HEK-293T cells are cultured in 10-cm plates at a high cell density of 375,000 cells/ml (3 x 10^6^ cells/8 ml) twenty-four hours prior to transfection. Cells are maintained using aseptic technique and are used in pharmacological assays within 5-30 passages.

### 2.2 Animals

C57Bl/6 mice are maintained in a vivarium provided by the Department of Laboratory Animal Medicine of Northeastern University. All experimental procedures performed on mice are in accordance with protocols approved by the Institutional Animal Care and Use Committee at Northeastern University.

### 2.3 Compounds & Plasmid Constructs

All CB1R ligands used in this study are purchased or acquired from Cayman Chemical (Ann Arbor, Michigan) or NIDA Drug Supply Program (Rockville, Maryland). The ligands are dissolved in DMSO to a stock concentration of 10 mM. All DNA constructs are generated in pcDNA3 plasmid vectors. Alanine substitutions of *CNR1*-containing plasmid constructs were generated using the Quickchange Site-Directed Mutagenesis Kit (Agilent) and the following primers:

F170A forward primer 5’ – TGGGGAGTGTCATTGCTGTCTACAGCTTCAT – 3’,

reverse primer 5’ – ATGAAGCTGTAGACAGCAATGACACTCCCCA – 3’;

S173A forward primer 5’ – TCATTTTTGTCTACGCCTTCATTGACTTCCA – 3’

reverse primer 5’ – TGGAAGTCAATGAAGGCGTAGACAAAAATGA 3’

F174A forward primer 5’ – ACAGCTTCATTGACGCCCACGTGTTCCACC – 3’,

reverse primer 5’ – GGTGGAACACGTGGGCGTCAATGAAGCTGT – 3’;

F177A forward primer 5’ – TTTTTGTCTACAGCGCCATTGACTTCCACG – 3’,

reverse primer 5’ – CGTGGAAGTCAATGGCGCTGTAGACAAAAA – 3’;

H178A forward primer 5’ – GCTTCATTGACTTCGCCGTGTTCCACCGCA – 3’,

reverse primer 5’ – TGCGGTGGAACACGGCGAAGTCAATGAAGC – 3’.

### 2.4 Bioluminescence Resonance Energy Transfer (BRET) receptor function assays

#### 2.4.1 General BRET methodology

BRET assays described below contain variations in DNA constructs transfected and in method of experimentation. Consistent with all experiments is a pre-incubation of 5-15 µg DNA (per 10-cm plate, DNA amounts vary by experiment) with 30 µg/cell plate linear polyethyleneimine (PEI) in serum-free DMEM twenty minutes before addition to cells. After overnight treatment, media is fully replaced with fresh, supplemented DMEM. After 48 hours of transfection, cells are washed with phosphate-buffered saline (PBS), harvested, and resuspended in PBS containing 0.1% glucose, 200 µM sodium bisulfite and 1 mg/mL fatty acid-free bovine serum albumin (BSA), which serves as a drug stabilizer. Cells are evenly distributed amongst wells in white flat-bottom 96 well plates. On the day of experiment, serially diluted drugs are transferred to cells three minutes after 5 µM coelenterazine H incubation. Luminescence and fluorescence values are recorded 10 minutes after drug treatment in a PheraStar FSX plate reader (bioluminescence at 480 nm, fluorescence at 530 nm). BRET ratio is calculated based on the measurement of fluorescence divided by that of luminescence. BRET ratios are then normalized to the basal BRET ratio calculated by the non-linear regression generated by GraphPad Prism 10.

#### 2.4.2 G-protein/β-arrestin engagement

Plasmid DNA in engagement BRET assays have been reported [31] as follows: CB1R tagged with *Renilla* luciferase 8 (CB1R-RLuc), Gα_i1_ tagged with Venus (Gα_i1_V), Gβ_1_, and Gγ_2_. For β-arrestin engagement, β-arrestin-2 tagged with Venus (βArr2V) and G-protein coupled receptor kinase 2 (GRK2) are used.

#### 2.4.3 G-protein activation

Plasmid DNA in activation BRET assays have been reported [35, 36] as follows: Untagged CB1R, Gα tagged with *Renilla* luciferase 8 (Gα_i1_-RLuc, Gα_i2_-RLuc, Gα_i3_-RLuc, Gα_oA_-RLuc, Gα_oB_-RLuc, or Gα_z_-RLuc), Gβ_1_, and Gγ_2_ tagged with Venus (Gγ_2_V).

### 2.5 Ex vivo slice electrophysiology

Young adult mice (8-20 weeks old) anesthetized with isoflurane were intracardially perfused with slicing artificial cerebral spinal fluid (slicing ACSF) containing (in mM): 92 NMDG, 20 HEPES, 25 glucose, 30 NaHCO3, 1.2 NaH2PO4, 2.5 KCl, 5 sodium ascorbate, 3 sodium pyruvate, 2 thiourea, 10 MgSO4, 0.5 CaCl2, 300–310 mOsm, at pH 7.3–7.4. After decapitation, the mouse brain is fixed to a stage, submerged in cold slicing ACSF saturated with 95% O2 and 5% CO2 (carbogen), and sectioned in a para-sagittal manner at 300 µm using a vibratome. Brain slices are transferred to 37°C slicing ACSF for seven minutes, then transferred to room temperature holding ACSF saturated with carbogen containing (in mM): 92 NaCl, 20 HEPES, 25 glucose, 30 NaHCO3, 1.2 NaH2PO4, 2.5 KCl, 5 sodium ascorbate, 3 sodium pyruvate, 2 thiourea, 1 MgSO4, 2 CaCl2, 300–310 mOsm, at pH 7.3–7.4. Slices are maintained in holding ACSF for 1.5 to 2 hours prior to experimentation. Brain slices are transferred into a recording chamber filled with recording ACSF that contains (in mM): 124 NaCl, 2.5 KCl, 1.25 NaH2PO4, 1 MgCl2, 26 NaHCO3, 11 glucose, 2.4 CaCl2, 0.05 caffeine, 300–310 mOsm, at pH 7.3–7.4. The recording ACSF is saturated with carbogen, maintained at 31-32°C, and perfused at 2 ml/min using a peristaltic pump. Caffeine, an adenosine A1 receptor antagonist, is used to reduce the effects of endogenous adenosine on observing CB1R function [37]. SCRAs are dissolved in ACSF containing 0.1% DMSO and 0.2-0.5% BSA, as SCRA lipophilicity can affect assay sensitivity [38]. Brain slices rest in perfused recording ACSF for 20 minutes before recording. A borosilicate recording electrode filled with recording ACSF is placed in the CA1 region of the hippocampus within the stratum radiatum, while a bipolar stimulating wire is placed in the CA3a region of the hippocampus. CA3 stimulation occurred at a frequency of 0.033 Hz, and were recorded using a MultiClamp 700b amplifier (10 kHz low-pass Bessel filter), digitized with a Digidata 1550b (20 kHz digitization), and visualized on pClamp software. Current amplitude is set at approximately 25-30% of max output, determined by generating an input/output curve. Baseline is recorded for 10 min before a 60-min drug perfusion.

### 2.6 Data Analysis

All data was processed and analyzed in GraphPad Prism 10 (San Diego, California). Data points were transformed to individual BRET ratio values and further normalized to the minimal and maximal responses by a reference full agonist CP55940 at 10 min as 0% and 100% respectively within each respective transducer. E_max_ and pEC_50_ parameters were obtained from the non-linear fit of normalized transformed data and multi-comparison data analyses at the 10 min time point were conducted. Based on the extrapolated curves, E_max_ and pEC_50_ are determined. E_max_ above 120% relative to the E_max_ of CP55,940 is interpreted as superagonist behavior. E_max_ on a non-plateaued curve and pEC_50_ less than 5 (i.e., 10 µM) are still reported in the tables. In those cases, E_max_ is reported as the efficacy at the highest concentration observed (10 µM). To evaluate whether SCRAs exhibited G protein subunit signaling bias, bias factors were calculated as reported previously [39]. This method yields bias factors similar to the operational model. Briefly, Δlog(E_max_/EC_50_) value for each transducer was calculated by subtracting the log(E_max_/EC_50_) value of the agonist by that of the reference compound. ΔΔlog(E_max_/EC_50_) is determined by subtraction of Δlog(E_max_/EC_50_) between two transducers. The comparison between the two transducers shows the bias direction of a certain ligand towards one of the two transducers. Throughout the experiments, a triplicate of at least four independent experiments was performed per each condition. Statistical grouped analyses on E_max_ and pEC_50_ were conducted using one-way ANOVA with a post-hoc Dunnett test for multiple comparisons.

For extracellular recordings, the fEPSP slope was determined in Clampfit from a 1.5-2 ms range in the rise phase post-stimulation. Recordings were averaged in 2-min bins and inhibition was determined as the fEPSP slope percentage as compared to the baseline fEPSP slope. Max inhibition was determined by averaging the fEPSP slope of the last six min of the drug prefusion. Tau was obtained from the non-linear regression of the time course. Statistical significance of max inhibition and tau was determined using one-way ANOVA with a post-hoc Dunnett test for multiple comparisons.

## 3. Results

### 3.1 L-tert-leucinate head moiety leads to high efficacy in SCRA derivatives

After recently investigating the drastic efficacy differences between l-valinate and l-*tert*-leucinate (*i.e.*, MMB- and MDMB-) head moieties of 5-fluoropenylindoles [31], we expanded our investigation on core and tail moieties in combination to the head moiety. Clinical studies have reported on the severe impact of indazole-based SCRA abuse, including death from overdose [3, 6, 40, 41]. Furthermore, original derivatives patented by Pfizer, as well as the drug responsible for the 2016 Brooklyn outbreak, carried a 4-fluorobenzyl tail moiety [6]. Therefore, we selected 5F-MMB-PICA (M-PC), 5F-MDMB-PICA (D-PC), 5F-MMB-PINACA (M-PN), 5F-MDMB-PINACA (D-PN), MMB-FUBICA (M-FC), MDMB-FUBICA (D-FC), MMB-FUBINACA (M-FN), MDMB-FUBINACA (D-FN) for our study panel to develop a SAR of these moieties for CB1R efficacy, thus investigating superagonism (Supp. Fig. 1).

We first measured the engagement of Gα_i1_ and β-arrestin-2 to CB1R as they represent two major signaling modalities, using an “engagement” mode of BRET where the receptor and Gα-protein or β-arrestin are fused with luciferase and fluorescent proteins, respectively (Fig. 1A, 1C). Both M-PC and D-PC displayed superagonism (i.e., above 120%), although M-PC is ∼20x less potent than D-PC in Gα_i1_ engagement (Fig. 1A). We see a separation in both efficacy and potency for the 4-fluorobenzyl analogues M-FC and D-FC (Fig. 1B). Interestingly, the difference in efficacy between MMB- and MDMB-compounds is negligible amongst indazole SCRAs (*i.e.*, M-PN, D-PN, M-FN, D-FN), as all confer superagonism, although there is still a noticeable potency shift (∼6-fold) for the MDMB-series (Fig. 1A-B, Table 1). Within the β-arrestin-2 engagement assay, seven of the eight SCRAs are more efficacious than the reference compound CP55,940, with only M-FC acting as a full agonist (Fig. 1C-D, Table 1). Although an insignificant difference for Gα_i1_ engagement, the 5-fluoropentyl SCRAs were all approximately 20-75% greater in efficacy than their 4-fluorobenzyl analogues (Fig. 1C-D, Table 1).

**Figure 1:**
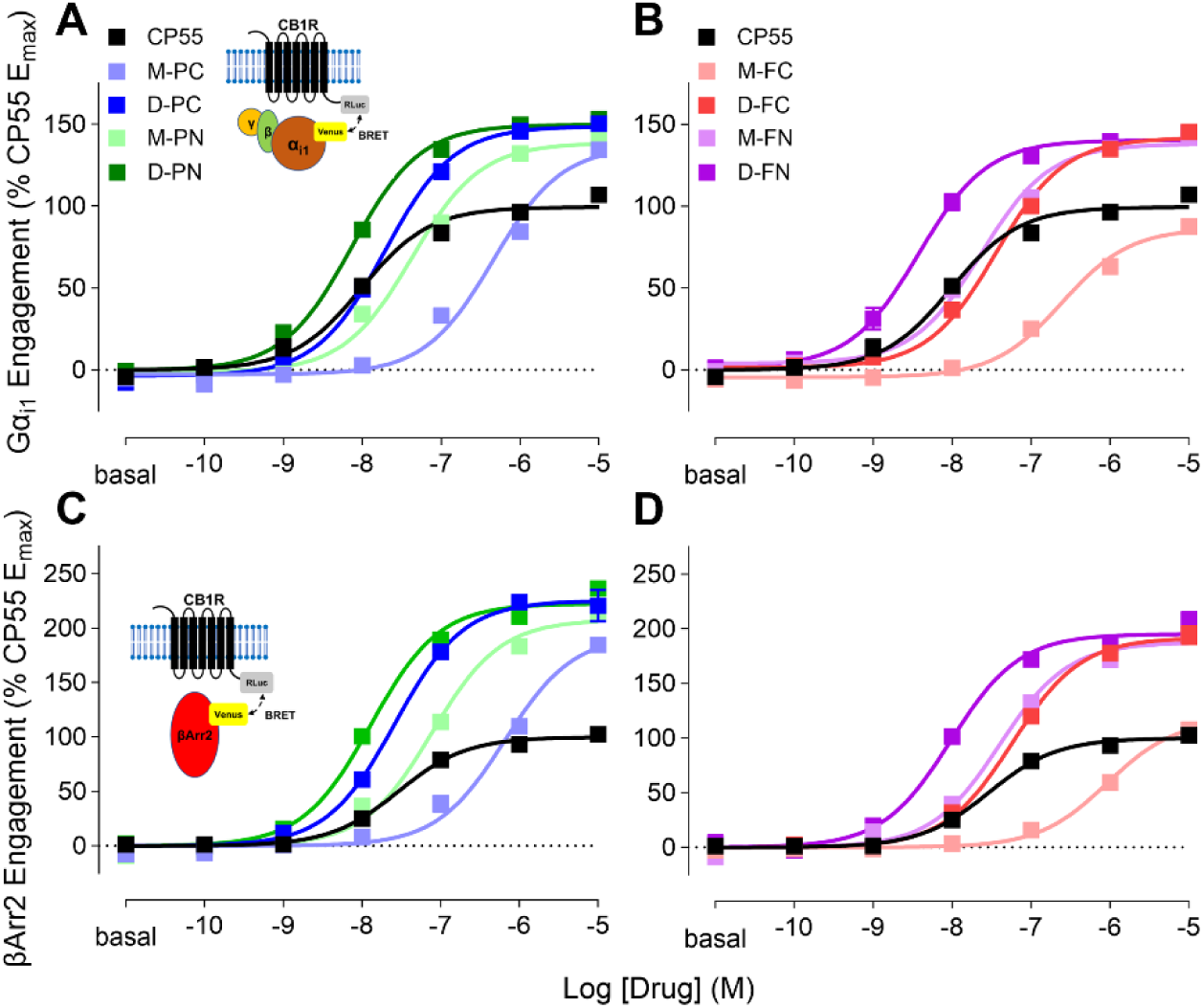
SCRA-induced Gα_i1_ and β-arrestin-2 engagement to CB1R. Drug-induced BRET between CB1R-RLuc and Gα_i1_-Venus (A,B) or βArr2-Venus (C,D) are shown. Concentration-response curves of SCRA-induced BRET are plotted as a percentage of CP55 E_max_ (A,B, Gα_i1_; C,D, βArr2) and organized by tail moiety (A,C 5-fluoropentyl; B,D 4-fluorobenzyl). Data are presented as means ± SEM of n=4 experiments.

**Table 1:** Pharmacological comparison of SCRAs in CB1R-transducer engagement BRET.

|  |  |  |  |  |  |  |
| --- | --- | --- | --- | --- | --- | --- |
|  | Gα<sub>i1</sub> |  | βArr2 |  | Bias Factor |  |
|  | E<sub>max</sub> (% CP55) | pEC<sub>50</sub> (M) | E<sub>max</sub> (% CP55) | pEC<sub>50</sub> (M) | ΔΔlog | Gα<sub>i1</sub> - βArr2 |
| **CP55** | 100.0 ± 1.75 | 8.01 ± 0.05 | 100.0 ± 1.59 | 7.54 ± 0.10 |  |  |
| **M-PC** | 136.5 ± 3.97* | 6.34 ± 0.06* | 193.7 ± 5.71* | 6.19 ± 0.05* | **M-PC - CP55** | -0.47 |
| **D-PC** | 148.6 ± 1.71* | 7.71 ± 0.03* | 225.1 ± 4.17* | 7.58 ± 0.06 | **D-PC - CP55** | -0.52 |
| **M-PN** | 138.5 ± 2.79* | 7.37 ± 0.06* | 207.6 ± 3.92* | 7.11 ± 0.06* | **M-PN - CP55** | -0.39 |
| **D-PN** | 149.6 ± 2.07* | 8.14 ± 0.04 | 222.5 ± 2.74* | 7.89 ± 0.09* | **D-PN - CP55** | -0.40 |
| **M-FC** | 86.5 ± 3.42* | 6.61 ± 0.09* | 117.3 ± 4.60* | 6.04 ± 0.13* | **M-FC - CP55** | -0.03 |
| **D-FC** | 131.0 ± 2.40* | 7.44 ± 0.06* | 191.9 ± 3.31* | 7.23 ± 0.05* | **D-FC - CP55** | -0.40 |
| **M-FN** | 137.8 ± 2.45* | 7.63 ± 0.05* | 187.5 ± 2.98* | 7.40 ± 0.08 | **M-FN - CP55** | -0.37 |
| **D-FN** | 140.2 ± 2.50* | 8.42 ± 0.06* | 195.2 ± 3.29* | 8.02 ± 0.08* | **D-FN - CP55** | -0.21 |

Data analyzed by one-way ANOVA followed by post-hoc Dunnett test. Data presented as means ± SEM. Significance of \* represents p<0.05 compared to CP55 E<sub>max</sub> and pEC<sub>50</sub> values for respective transducer.

### 3.2 SCRA superagonism is observed across activation of most Gα_i/o_ subtypes

Numerous studies have reported on SCRA-mediated CB1R activities without identifying the involvement of specific Gα_i/o_ subtypes. Concurrently, there are limited findings on SCRA agonism amongst the six inhibitory G proteins (Gα_i1_, Gα_i2_, Gα_i3_, Gα_oA_, Gα_oB_, Gα_z_). Therefore, we studied the extent of SCRA superagonism across all six Gα_i/o_ subtypes, as well as the contribution of key moieties of these SCRAs in the functional selectivity.

To investigate this, we use an “activation” mode of BRET to measure the G-protein heterotrimer dissociation via untagged CB1R (Fig. 2A, Table 2) [36, 42, 43]. In general, MDMB-compounds are more potent than MMB-compounds. Amongst indole SCRAs, there was a noticeable increase in both efficacy and potency for MDMB series compared to MMB series (Fig. 2B-G), but only an efficacy difference (>20%) for M-FC and D-FC in line with the Gα_i1_ “engagement” assay results (Fig. 1A-B). M-FC showed the lowest efficacy and potency for Gα_i_ subtypes; in the latter case, sharing the least potency with M-PC. For indazoles, MDMB-compounds are more potent than MMB-compounds, but there is no clear difference in efficacy in general. In Gα_oA_, we found that D-PN was the single instance of an MDMB-indazole displaying noticeably greater efficacy (∼50%) than its MMB-analogue (Fig. 2D, Table 2). Interestingly, a general high efficacy trend among all SCRAs tested (except for D-FC) was observed for Gα_i3_ (>150%), Gα_oA_ (>125%), and Gα_z_ (>124%).

**Figure 2:**
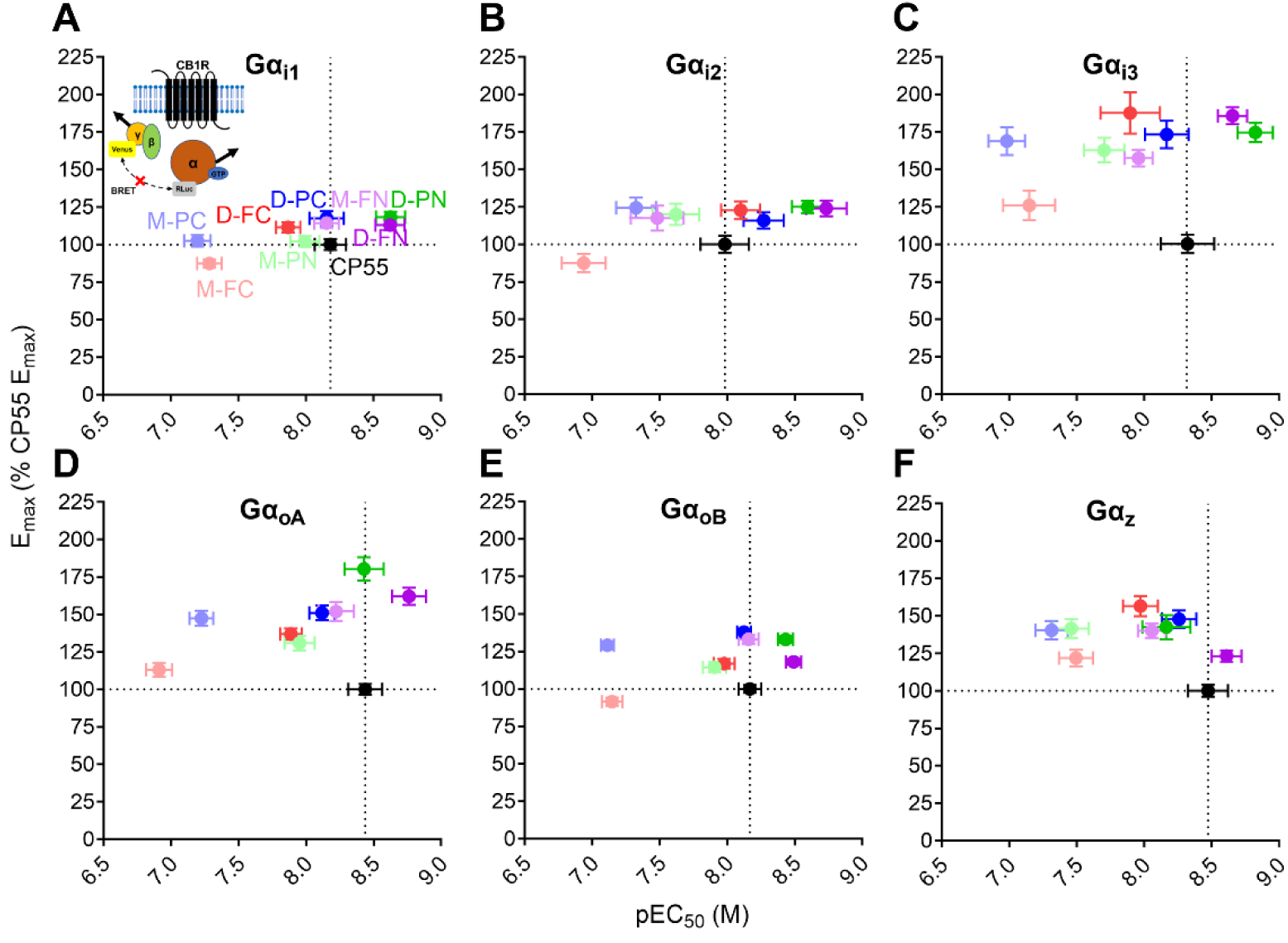
SCRA-activated Gα_i/o_ subtype protein dissociation BRET assay. Cartoon illustrating BRET interaction between CB1R and Gα-RLuc and Gγ_2_-Venus with untagged CB1R (A, insert). SCRA-induced BRET efficacy, as a percentage of CP55 E_max_, and pEC50 were plotted for Gα_i/o_ subtype transducers: Gα_i1_ (A), Gα_i2_ (B), Gα_i3_ (C), Gα_oA_ (D), Gα_oB_ (E), and Gα_z_ (F). Data presented as means ± SEM of n≥4 experiments. The full concentration-response curves can be found in Supplementary Figure 2.

**Table 2:**
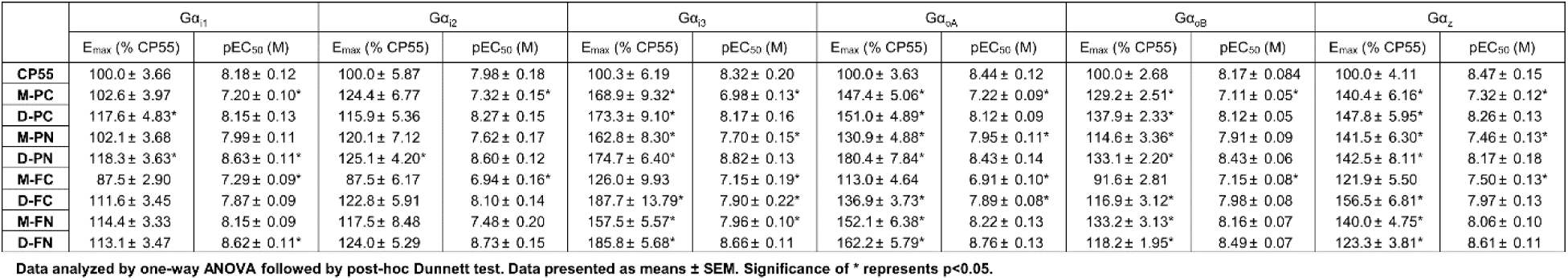
Pharmacological comparlsori of SCR.As In G protein dissociation BRET.

We further analyzed Gα_i/o_ subtype activation by estimating each compound’s functional selectivity by calculating bias factors. Here, we determined the Δlog(E_max_/EC_50_) of each SCRA to compare two different transducers then taking the difference between the Δ values of a particular SCRA and the reference agonist CP55,940, obtaining ΔΔlog(E_max_/EC_50_) (Table 3). Here we found that none of the compounds had a ΔΔlog value outside the range of -1 to 1 (i.e., 10-fold separation between two Gα_i/o_ subtypes relative to the reference CP55,940), thus suggesting the tested SCRAs are not significantly biased towards any specific pathway.

**Table 3:**
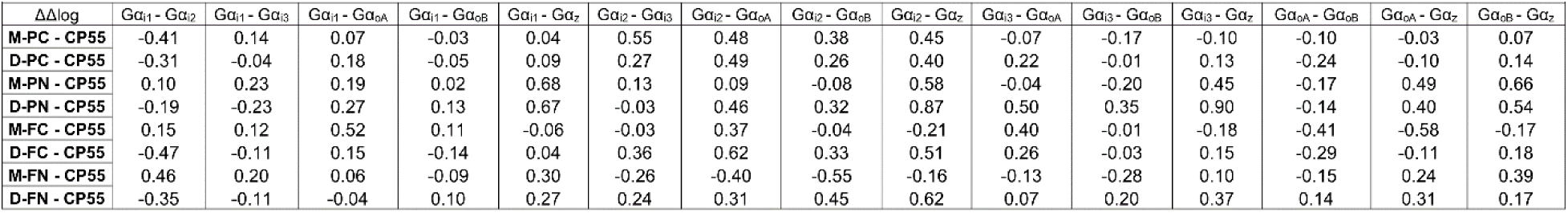
SCRA-mediated activation bias of Gα_i/o_ subtypes.

### 3.3 F170, F177, and H178 are key residues in SCRA activation, not exclusive to l-tert-leucinate

After having established the consistent increase in efficacy and potency for l-*tert*-leucinate (*i.e.*, MDMB-) containing SCRAs, we investigated residues within the binding pocket essential for conferring superagonism. Recent research has found that extracellular transmembrane domain 2 (TM2) residues are implicated in CB1R activation as they rotate inwards in the presence of a full agonist [32, 33]. Indeed, we previously reported on the conformational stability of TM2 being associated with 5F-MDMB-PICA superagonism [31]. Here, we expanded our investigation by generating alanine substituted mutants along the TM2 residues in order to isolate the essential residues implicated in CB1R superagonism.

To that end, we generated alanine-substituted CB1R constructs at F170, S173, F174 F177, and H178, intracellular to extracellular on TM2, and assessed SCRA activity in these mutant constructs through Gα_i1_ and β-arrestin-2 engagement BRET assays (Fig. 3-4). We selected the 5-fluoropentyl SCRAs (M-PC, D-PC, M-PN, D-PN) for this study, as these were the more efficacious compounds respective to their 4-fluorobenzyl analogs (Fig. 3A). By normalizing the E_max_ of CP55,940 for the wildtype CB1R as 100% efficacy, mutational effects on the SCRAs were studied.

**Figure 3:**
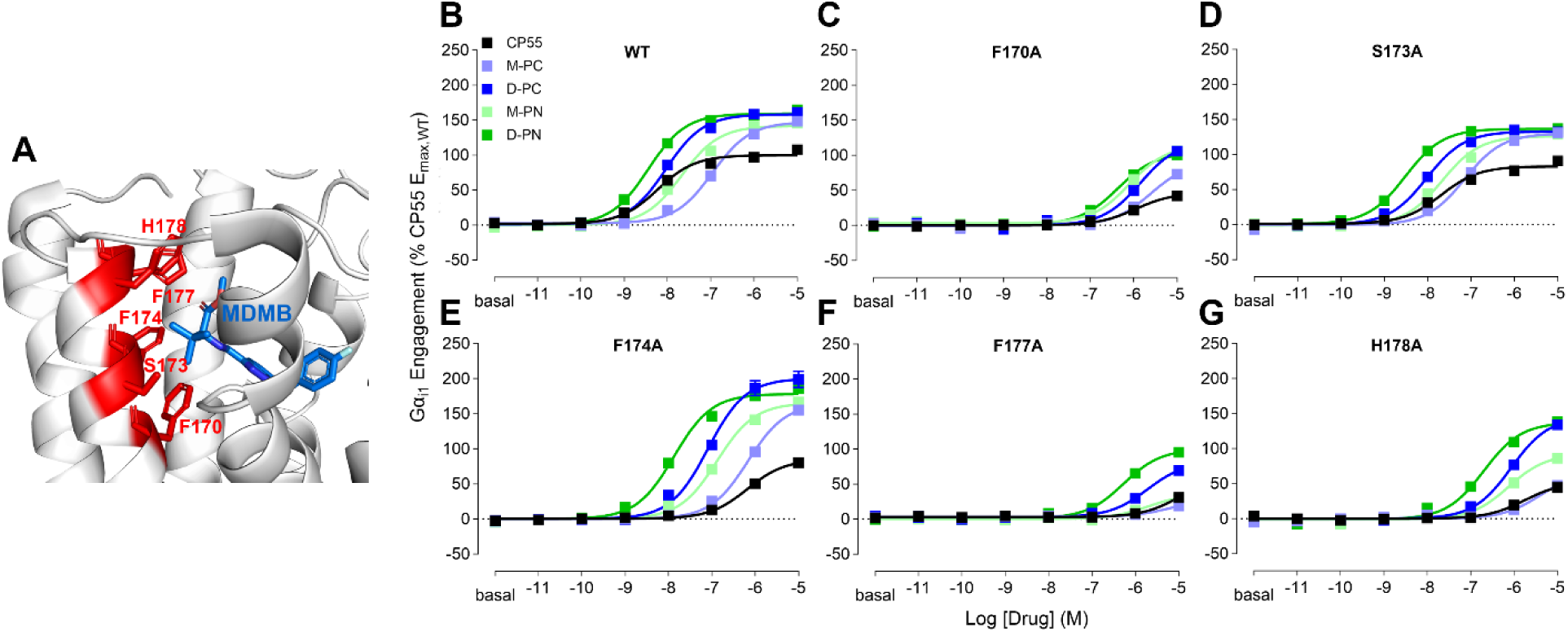
CB1R-TM2 mutational analysis of G-protein engagement. A close-up view of TM2-MDMB-FUBINACA (PDB:6N4B) with residues (red) within 4 Å of the MDMB-head moiety (blue) using PyMOL 3.0 (Schrödinger) for visualization (A). Through BRET Gα_i1_ Engagement mode, M-PC, D-PC, M-PN, and D-PN are compared to CP55 in the presence of CB1R with the following TM2 mutations: Wildtype (B), F170A (C), S173A (D), F174A (E), F177A (F), H178A (G). Concentration-response curves of SCRA-induced BRET plotted as a percentage of CP55 E_max,WT_. Data presented as means ± SEM of n=4 experiments.

**Figure 4:**
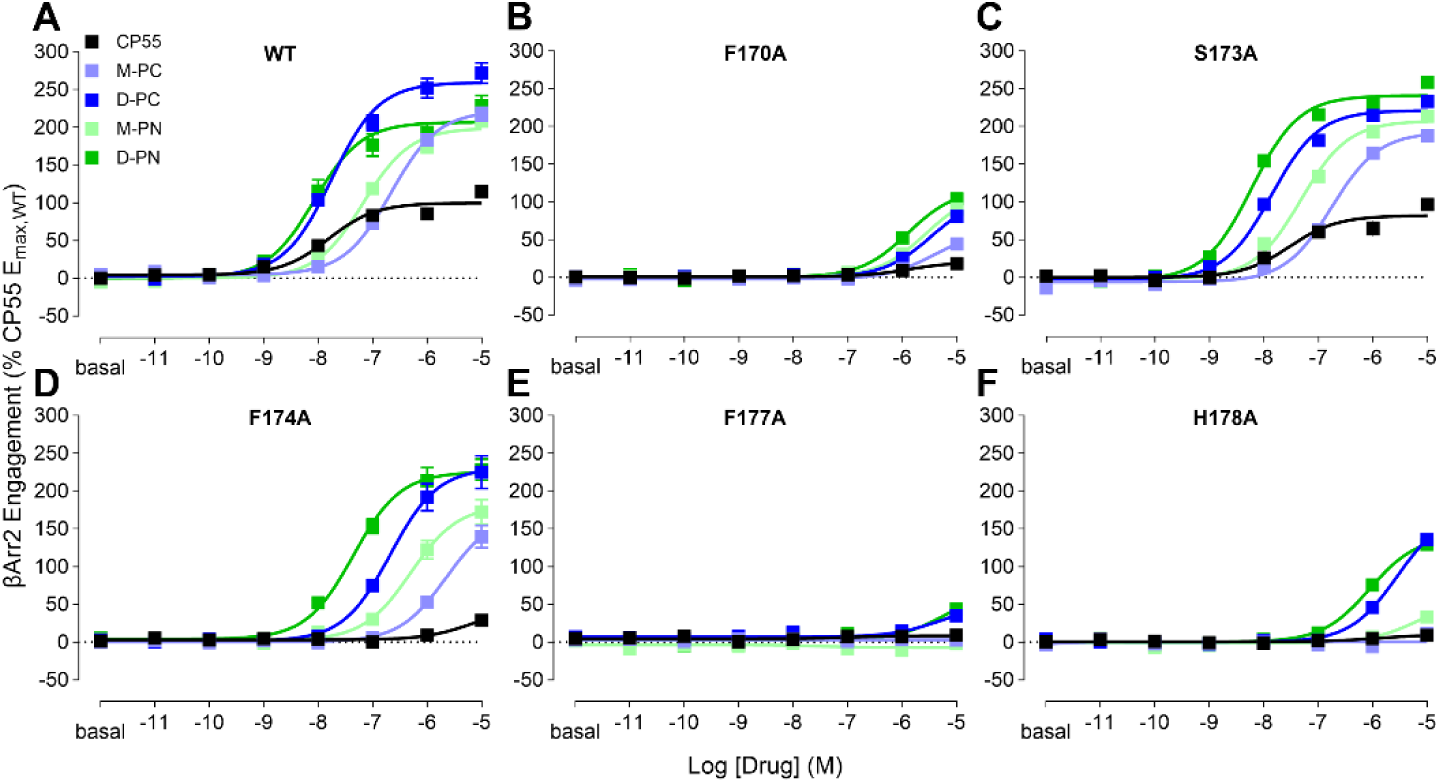
CB1R-TM2 mutational analysis of β-arrestin-2 engagement. Through BRET βArr2 Engagement mode, M-PC, D-PC, M-PN, and D-PN are compared to CP55 in the presence of CB1R with the following TM2 mutations: Wildtype (A), F170A (B), S173A (C), F174A (D), F177A (E), H178A (F). Concentration-response curves of SCRA-induced BRET plotted as a percentage of CP55 E_max,WT_. Data presented as means ± SEM of n=4 experiments.

In the Gα_i1_ engagement assay, the F170A substitution blunted CP55,940 efficacy, and reduced the efficacy and potency of all the SCRAs tested, although M-PC had a lower efficacy than the other three (Fig. 3C, Table 4). The S173A substitution produced no significant change to the efficacy and potencies of the drugs tested (Fig. 3D, Table 4). Although F174A reduced the potency of the agonists, we observed a slight increase in the efficacy of these SCRAs relative to CP55,940 activity in wildtype CB1R (Fig. 3E, Table 4). The F177A mutation nearly blunted the efficacy of CP55,940, M-PC and M-PN, but only partially for D-PC and less so D-PN (Fig. 3F, Table 4). Similarly to F177A, the H178A mutation reduced the efficacy of CP55,940 and M-PC to low partial agonism, M-PN to high partial agonism, whereas D-PC and D-PN conferred high agonism (Fig. 3G, Table 4).

**Table 4:** CB1R-TM2 Mutational An.alysls with SCRA Head Moiety (Gα_i1_ Engagement)

| Table 4: CB1R-TM2 Mutational Analysis with SCRA Head Moiety (G<sub>αi</sub> Engagement) |  |  |  |  |  |  |  |  |  |  |  |  |
| --- | --- | --- | --- | --- | --- | --- | --- | --- | --- | --- | --- | --- |
|  | WT |  | F170A |  | S173A |  | F174A |  | F177A |  | H178A |  |
|  | E<sub>max</sub> (% CP55,WT) | pEC<sub>50</sub> (M) | E<sub>max</sub> (% CP55,WT) | pEC<sub>50</sub> (M) | E<sub>max</sub> (% CP55,WT) | pEC<sub>50</sub> (M) | E<sub>max</sub> (% CP55,WT) | pEC<sub>50</sub> (M) | E<sub>max</sub> (% CP55,WT) | pEC<sub>50</sub> (M) | E<sub>max</sub> (% CP55,WT) | pEC<sub>50</sub> (M) |
| CP55 | 100.0 ± 1.31 | 8.23 ± 0.04 | 42.2 ± 3.78 | N.D. | 83.1 ± 1.62 | 7.70 ± 0.06 | 85.0 ± 3.41 | 6.18 ± 0.08* | 31.0 ± 2.17 | N.D. | 44.7 ± 3.33 | N.D. |
| M-PC | 147.4 ± 2.06* | 6.96 ± 0.03* | 72.9 ± 3.02 | N.D. | 130.1 ± 1.96* | 7.15 ± 0.04* | 164.9 ± 4.15* | 6.17 ± 0.05* | 18.8 ± 1.27 | N.D. | 47.5 ± 2.74 | N.D. |
| D-PC | 158.3 ± 1.78* | 8.04 ± 0.03 | 121.3 ± 4.59* | 5.84 ± 0.06* | 132.8 ± 1.15* | 8.02 ± 0.03 | 200.4 ± 4.99* | 7.09 ± 0.06* | 69.5 ± 2.89 | N.D. | 145.3 ± 3.83* | 6.06 ± 0.05* |
| M-PN | 141.8 ± 1.85* | 7.67 ± 0.04 | 114.7 ± 3.85 | 6.01 ± 0.06* | 126.3 ± 1.50* | 7.67 ± 0.03 | 165.3 ± 2.43* | 6.88 ± 0.03* | 29.9 ± 3.21 | N.D. | 92.5 ± 3.93 | 6.12 ± 0.08* |
| D-PN | 159.4 ± 1.91* | 8.45 ± 0.04 | 104.1 ± 3.34 | 6.37 ± 0.07* | 136.6 ± 1.14* | 8.52 ± 0.03 | 178.3 ± 1.73* | 7.87 ± 0.03 | 100.3 ± 2.89 | 6.27 ± 0.06* | 137.6 ± 2.75* | 6.70 ± 0.05* |

Data analyzed by one-way ANOVA followed by post-hoc Dunnett test. Data presented as means ± SEM. Significance of \* represents p<0.05 compared to CP55 E<sub>max</sub>, pEC<sub>50</sub> of the wildtype construct. For concentration-response curves with a low potency shift and/or low E<sub>max</sub>, the pEC<sub>50</sub> was not determined (N.D.) and the E<sub>max</sub> shown is the effect at the maximum concentration tested (10 μM).

When comparing these results to β-arrestin-2 engagement, we find an overall rightward potency shift across the mutant constructs (Table 5). In F170A and S173A, we see similar efficacy reduction that we observed in Gα_i1_ engagement (Fig. 4B-C). Although efficacy increased in CB1R F174A Gα_i1_ engagement, SCRA efficacy was slightly reduced, relative to wildtype CB1R activity, in CB1R F174A β-arrestin-2 engagement. However, CP55,940 showed much higher efficacy reduction (71.5%) relative to wildtype, suggesting a differential residue-ligand interaction at F174 (Fig. 4D, Table 5). Interestingly, all drug response was flattened in CB1R F177A β-arrestin-2 engagement, where only l-*tert*-leucinate compounds had >30% efficacy (Fig. 4E, Table 5). In the H178A mutation results, we observed high efficacy in l-tert-leucinate compounds, and less than 35% efficacy in all other compounds (Fig. 4F, Table 5).

**Table 5:** CB1R-TM2 Mutational Analysis withSCRA Head Moiety (βArr2Engagement)

|  |  |  |  |  |  |  |  |  |  |  |  |  |
| --- | --- | --- | --- | --- | --- | --- | --- | --- | --- | --- | --- | --- |
|  | WT |  | F170A |  | S173A |  | F174A |  | F177A |  | H178A |  |
|  | E<sub>max</sub> (% CP55,WT) | pEC<sub>50</sub> (M) | E<sub>max</sub> (% CP55,WT) | pEC<sub>50</sub> (M) | E<sub>max</sub> (% CP55,WT) | pEC<sub>50</sub> (M) | E<sub>max</sub> (% CP55,WT) | pEC<sub>50</sub> (M) | E<sub>max</sub> (% CP55,WT) | pEC<sub>50</sub> (M) | E<sub>max</sub> (% CP55,WT) | pEC<sub>50</sub> (M) |
| **CP55** | 100.0 ± 2.76* | 7.82 ± 0.08 | 18.3 ± 1.94 | N.D. | 81.6 ± 3.90 | 7.53 ± 0.14 | 28.5 ± 3.51 | N.D. | 8.82 ± 1.84 | N.D. | 8.61 ± 4.39 | N.D. |
| **M-PC** | 221.9 ± 4.47* | 6.66 ± 0.05 | 43.9 ± 6.77 | N.D. | 192.4 ± 7.79* | 6.76 ± 0.09 | 170.7 ± 12.36* | 5.65 ± 0.11 | 2.71 ± 2.56 | N.D. | 11.3 ± 2.10 | N.D. |
| **D-PC** | 259.3 ± 5.39* | 7.76 ± 0.06 | 80.7 ± 7.77 | N.D. | 221.1 ± 7.06* | 7.86 ± 0.09 | 230.7 ± 9.67* | 6.68 ± 0.10 | 34.9 ± 2.66 | N.D. | 135.6 ± 6.39 | N.D. |
| **M-PN** | 199.0 ± 4.54* | 7.17 ± 0.06 | 90.4 ± 6.60 | N.D. | 207.3 ± 8.10* | 7.32 ± 0.10 | 181.2 ± 8.75* | 6.30 ± 0.10 | -2.0 ± 3.44 | N.D. | 33.4 ± 3.45 | N.D. |
| **D-PN** | 206.6 ± 6.56* | 8.07 ± 0.10 | 117.1 ± 6.06 | 5.9 ± 0.09 | 240.9 ± 8.86* | 8.23 ± 0.11 | 225.3 ± 6.71* | 7.356 ± 0.08 | 43.5 ± 5.29 | N.D. | 141.0 ± 4.79 | 6.05 ± 0.06 |

Data analyzed by one-way ANOVA followed by post-hoc Dunnett test. Data presented as means ± SEM. Significance of \* represents p<0.05 compared to CP55 E<sub>max</sub>, pEC<sub>50</sub> of the wildtype construct. For concentration-response curves with a low potency shift and/or low E<sub>max</sub>, the pEC<sub>50</sub> was not determined (N.D.) and the E<sub>max</sub> shown is the effect at the maximum concentration tested (10 μM).

### 3.4 L-tert-leucinate SCRAs confer high efficacy agonism in glutamate release inhibition in the hippocampus

To investigate the pharmacological profiles of SCRAs at a neurophysiological level, we evaluated the effects of l-*tert*-leucinate SCRAs through *ex vivo* brain slice electrophysiology. Specifically, CB1R-mediated suppression of neurotransmitter release was studied in the Schaffer collaterals of the hippocampus, as we have shown [31]. In the previous study, we reported that D-PC is significantly higher in efficacy compared to M-PC in inhibiting glutamate release in the hippocampus. Here, we evaluated the efficacy and kinetics of other l-*tert*-leucinate SCRAs on glutamate release.

We performed para-sagittal slicing of young adult mice brains to reveal the hippocampus and then perfused 1 µM, 300 nM, and 100 nM of SCRAs over the course of 60 minutes while recording field excitatory postsynaptic potentials (fEPSP) in the hippocampus (Fig. 5A, Supp. Fig. 4). l-valinate-containing M-PC was used as a reference SCRA against all l-*tert*-leucinate-containing (i.e., MDMB-) SCRAs. As expected, M-PC showed a low response, resulting in approximately 25% inhibition of the fEPSP slope. All four MDMB-compounds resulted in 40-55% inhibition, significantly greater than that of M-PC at 1 µM (Fig. 5B). When compared at lower concentrations, only 300 nM D-FC had a significantly lower inhibition than 1 µM (Fig. 6B). We also compared the kinetics of efficacy increase via CB1R over time with tau values of these compounds. We found that the tau for 1 µM D-PC and D-PN were under 10 minutes, whereas 1 µM D-FC and D-FN were ∼10-20 minutes greater than that of M-PC, although these values were not determined to be significant (Fig. 5C). Although reaching significantly higher efficacy than M-PC, MDMB-SCRAs showed differential kinetics, specifically, slower response with 4-fluorobenzyl tail ones.

**Figure 5:**
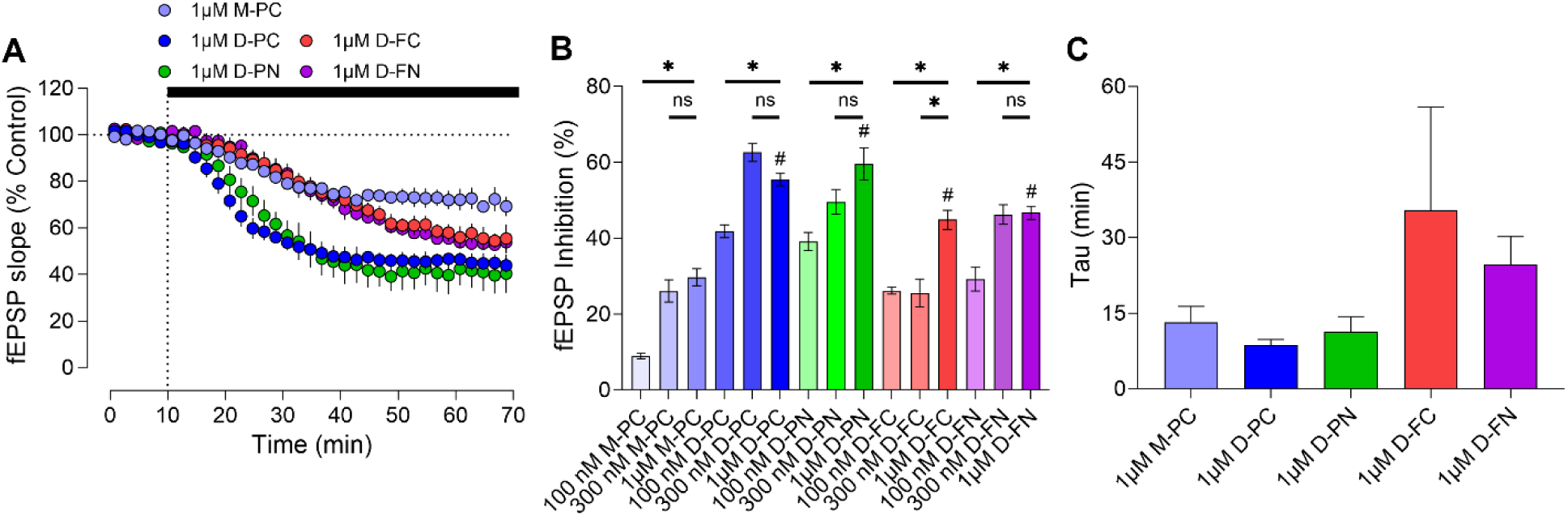
Extracellular field recordings in the CA1 hippocampus after perfusion of L-*tert*-leucinate SCRAs. Field extracellular postsynaptic potentials (fEPSPs) were recorded every 30 seconds in the CA1 hippocampus during SCRA perfusion. After 10 minutes of baseline, SCRAs were perfused for 60 minutes (A). Maximum inhibition was calculated from the average of the last six minutes of 1 µM, 300 nM, 100 nM SCRA perfusion (B). The time decay constant tau was calculated from the fitted curve of the timecourse recording (C). (N≥4 mice; n≥4 recordings; means ± SEM; p<0.05 compared to respective 1 µM SCRA = *, p<0.05 compared 1 µM M-PC = #.)

## 4. Discussion

Previous reports by others assessed a number of SCRAs used here as “high efficacy” agonists [28, 44, 45]. By using the BRET “engagement” mode, we can better address issues associated with signal amplification and receptor reserves, which can result in falsely high efficacy and potency activity in downstream functional assays [30, 46], whereas the engagement mode detect direct coupling between the receptor and transducer. Contrary to our previous results, M-PC displays superagonism for both Gαi1 and β-arrestin-2 engagement modes of BRET assays [26]. This likely resulted from adding 1 mg/mL BSA to the assay buffer as a carrier protein to these highly lipophilic compounds for their solubility and stability, as used in previous studies [21,22]. Nonetheless, D-PC shows remarkably higher potency compared to M-PC, consistent with our previous finding [31]. Indazoles, regardless of head or tail moiety within the structure, drastically increases SCRA efficacy and potency, the latter of which has been well reported for indazole SCRA derivatives [25, 26]. The superagonist activity of D-PC, D-PN, and D-FN, in both Gα_i/o_ and β-arrestin engagement, likely contribute significantly to the severity of the adverse effects, as cannabis use, thus moderate efficacy of THC on CB1R, rarely causes the severe adverse effects and toxicities [10]. The increased efficacy in β-arrestin-2 engagement may be associated with increased tolerance found in SCRA use as β-arrestins mediate CB1R internalization [47, 48]. Indeed, SCRAs have been reported to induce rapid CB1R internalization through β-arrestin [26, 29]. These studies as well as our findings indicate that high efficacy in β-arrestin likely plays a role in adverse effects reported in human use.

Through G protein activation BRET assays, we determined that the panel of tested SCRAs displays superagonism across Gα_i/o_ subtypes. This finding is crucial as most previous reports have not focused on SCRA characterization of all Gα_i/o_ subtypes as biased responses in different organs and cell-types can be implicated through diverse Gα_i/o_ subtype expression. Indeed, the Gα_i1-3_ are ubiquitously expressed throughout the body and are the dominant Gα_i/o_ subtypes in the periphery, as the Gα_oA-B_ and Gα_z_ are the dominant proteins in the CNS. SCRA-induced superagonism across these subtypes may suggest a connection to the wide-range of adverse effects from SCRA use. We attempted to reveal any potential bias across head, core, and tail moieties by calculating ΔΔΔlog values from ΔΔlog of differing moieties (data not shown). However, we did not identify any significant bias beyond the -1 or 1 threshold, thus none of the moieties cause bias within Gα_i/o_ subtypes. Other clinically reported SCRAs have also been found relatively balanced using similar *in vitro* assays [26–28]. Since there are a multitude of combinatorial diversity among different head, core and tail moieties for SCRAs, characterization of signaling bias remains to be explored and incomplete at present. Regardless, we have determined that across Gα_/o_ subtypes, the panel of paired SCRA analogs at head, core and tail moieties display CB1R superagonism.

In the mutational study, we investigated the TM2 residues implicated in SCRA-mediated CB1R superagonism. Recent reports have indicated that H178 is implicated in the activation of the CB1R [32, 34]. We confirmed that to be the case as the alanine substitution drastically reduced the effect of CP55,940, M-PC, and M-PN in both engagement modes. This phenomenon implicates the ring structure and/or positive charge of histidine as essential in CB1R engagement to an agonist in general, beyond head moiety interaction in the aminoalkylindole agonists, warranting further investigation. We also found CP55,940 response was completely suppressed in F170A, F174A, and F177A, supporting results found recently in cAMP assays [34]. For the SCRAs, we expected that differences in the head moiety would lead to noticeable differences in activity in the more extracellular alanine substitutions in TM2, whereas the core moiety would be highlighted in the more intracellular substitutions. The alanine substitution at F170, the most intracellular residue tested, highlighted the difference in efficacy between MMB- and MDMB-compounds, but only for the indole compounds (M-PC, D-PC). Although M-PN has the less efficacious l-valinate head moiety, it remains as effective as D-PC and D-PN, suggesting an importance in the indazole stabilizing the active conformation. At only one residue down from the extracellular end H178 in TM2, F177A exhibited a clear separation in efficacy between MMB- and MDMB-compounds. Furthermore, this residue may be implicated in β-arrestin-2 bias, as even the l-*tert-* leucinate compounds showed significantly reduced efficacy, a distinct profile compared to that of F170 and H178. F174, in the middle of F170 and F177, seems necessary for CP55,940 activity, yet does not substantially impact SCRA efficacy and potency. To further characterize F174A, we tested the construct in Gα_oA_ activation and found that the potency was reduced yet the efficacy had dramatically increased (Supp. Fig. 2D, Supp. Table 1). We found that the basal net BRET ratio for F174A was much higher than WT in Gα_oA_ activation, indicating a lower constitutive receptor activity by the F174A mutation (Supp. Fig. 3C). However, the F174A construct remains capable of reaching levels of BRET on par with the wildtype at saturating agonist concentration, thus reaching the calculated high percentage of efficacy. We expected the head moiety of M-PC and D-PC to interact with S173 and, therefore, the S173A mutation to lower their efficacy [31]. Nonetheless, in our engagement assays, we did not observe a drastic efficacy decrease with S173A as in F170A and F177A. We did find an overall reduced efficacy in CB1R S173A Gα_oA_ activation, yet SCRA efficacy and potency relative to CP55,940 efficacy and potency was similar to those found in wildtype CB1R Gα_oA_ activation. (Supp. Fig. 2C, Supp. Table 1). Overall, S173 seems to have a smaller role in CB1R activity compared to the other TM2 residues tested. Further investigation of interacting residues beyond these TM2 residues will provide more insight into SCRA superagonism and key interactions with SCRA core and tail moieties as well.

In the field recording of hippocampal slices, we confirmed that l-*tert*-leucinate SCRAs confer a high degree of efficacy at a neurophysological level. Interestingly, M-PC confers a distinctly lower level of efficacy to D-PC in electrophysiology that we do not see in our BRET studies. That is, despite the addition of BSA, field recordings did not differ much from our previous results. However, when compared to BRET results with BSA addition, the significant efficacy difference between M-PC and D-PC is gone. Although all MDMB-compounds had much greater levels of inhibition, the tail moiety seemed to play some role in the kinetics of these drugs, given the higher tau for 4-fluorobenzyl (i.e., D-FC and D-FN) than 5-fluoropentyl compounds (i.e., M-PC, D-PC, and D-PN). This difference may be due to the higher lipophilicity of the benzyl ring in the tail moiety (e.g., cLogP values: M-PC – 3.9, D-PC – 4.3, D-FC – 4.7, D-PN – 4.2, D-FN – 4.6), potentially leading to reduced solubility and receptor access. Indeed, kinetic differences were not observed among MDMB-compounds in *in vitro* BRET assays as there is no tissue penetration involved for the receptor activity. Within the 4-fluorobenzyl SCRAs, we do see that the indole (D-FC) is less potent than D-FN, given the difference in efficacy at 300 nM. As such our electrophysiological recordings can be a viable method to distinguish ligand accessibility to native CB1Rs and evaluate their efficacy at neurons.

## 5. Conclusion

The iterative design of SCRAs in their modular components has led to an explosive number of substances, many manifested toxicities. In this study, we attempted to understand the involvement of TM2 in a SAR for SCRA-induced CB1R superagonism. Using structural pairwise comparison within head, core, and tail moieties, we detailed superagonism conferred by the l-*tert*-leucinate head moiety. We found that SCRAs, depending on their key moieties, can elicit superagonism through multiple Gα_i/o_ subtypes and β-arrestin, and that F170, F177, and H178 in TM2 are implicated in conferring such drug action. This pairwise comparison approach within ligand structure in conjunction with extensive transducer analysis may be useful for revealing specific activation profiles in the future SAR studies for novel SCRAs.

## CRediT authorship contribution statement

**Christopher Lucaj:** Conceptualization, Data curation, Formal analysis, Investigation, Methodology, Validation, Visualization, Writing – original draft, Writing – review & editing. **Soo Jung Oh:** Investigation, Methodology **Charlotte Pitha:** Investigation, Methodology **Jordan Davis:** Investigation, Methodology **Hideaki Yano:** Conceptualization, Data curation, Formal analysis, Project administration, Resources, Funding acquisition, Supervision, Writing – original draft, Writing – review & editing

## Acknowledgements and Funding

The work is supported by the Brain & Behavior Research Foundation Young Investigator Grant (H.Y.) and by the National Institute on Drug Abuse (T32DA055553 to C.L.) We would like to thank Dr. Carl Lupica and Dr. Alexander Hoffman for technical advice and discussion on brain slice electrophysiology.

---

**Supplementary Figure 1:**
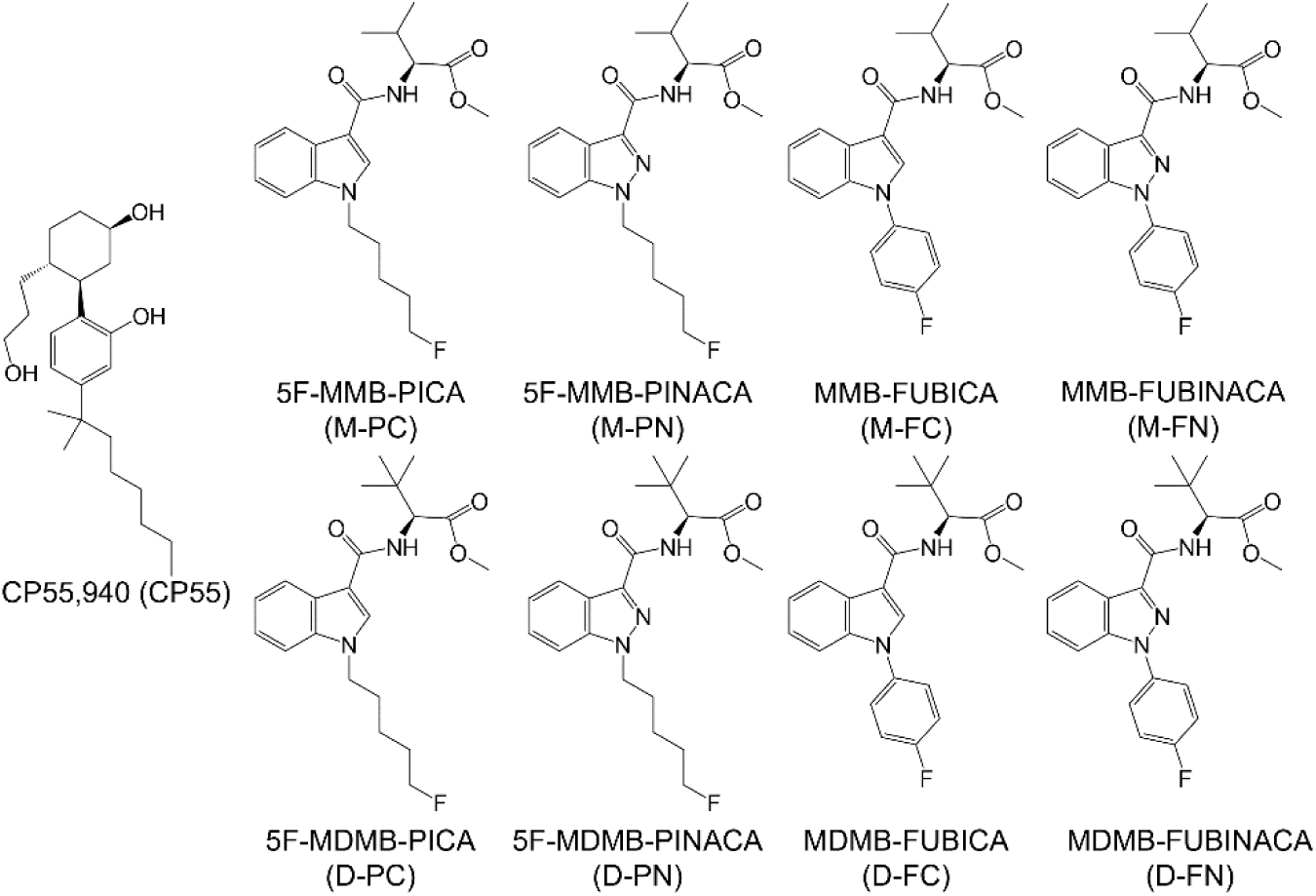
Structures of SCRAs used in this study. CP55,940 serves as the reference ligand for all experiments.

**Supplementary Figure 2:**
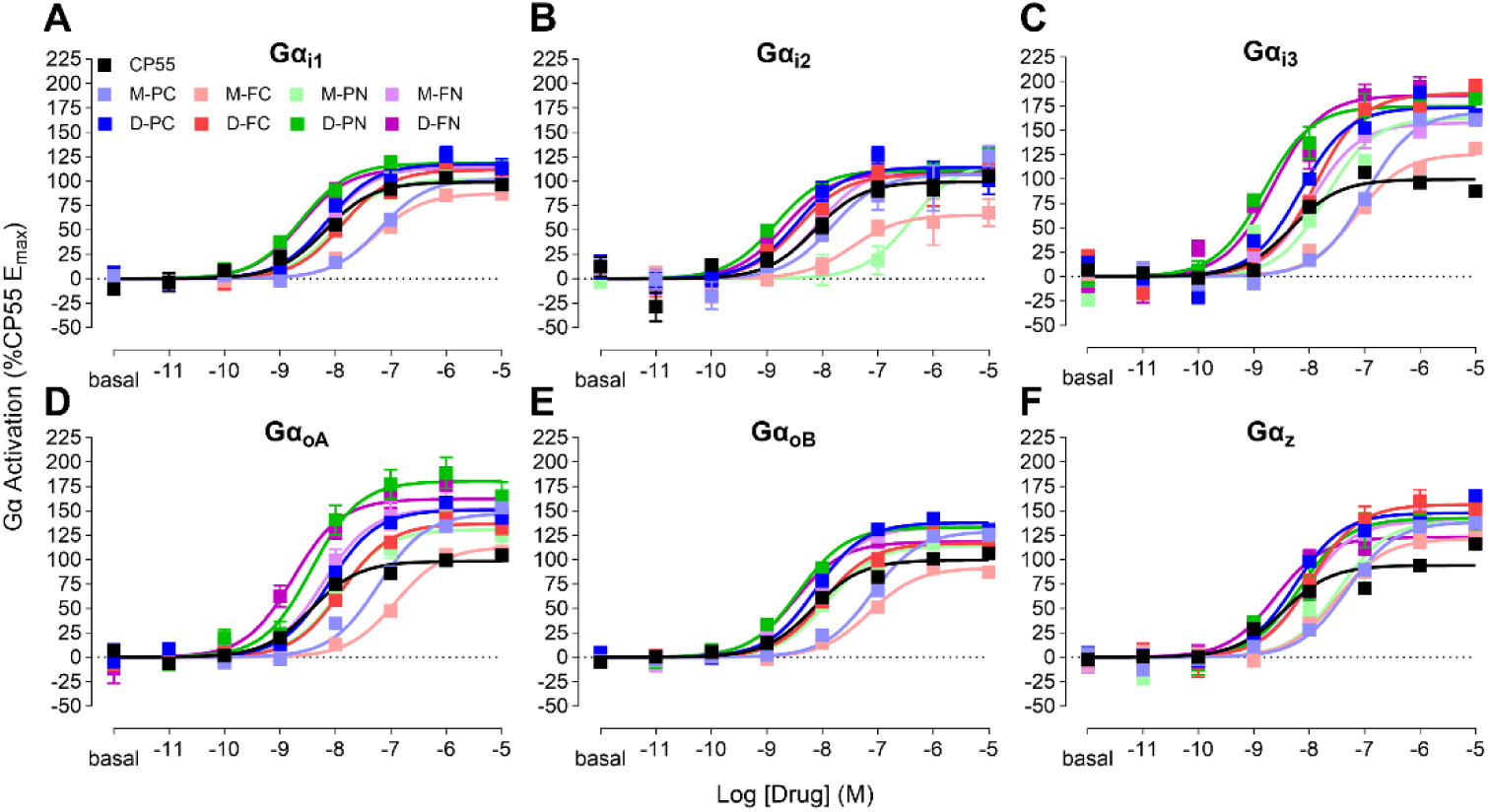
Full concentration-response curves of SCRA-mediated G-protein activation. Concentration-response curves of SCRA-induced BRET are plotted as a percentage of CP55 E_max_ for each transducer: Gα_i1_ (A), Gα_i2_ (B), Gα_i3_ (C), Gα_oA_ (D), Gα_oB_ (E), and Gα_z_ (F). Data are presented as means ± SEM of n≥4 experiments.

**Supplementary Figure 3:**
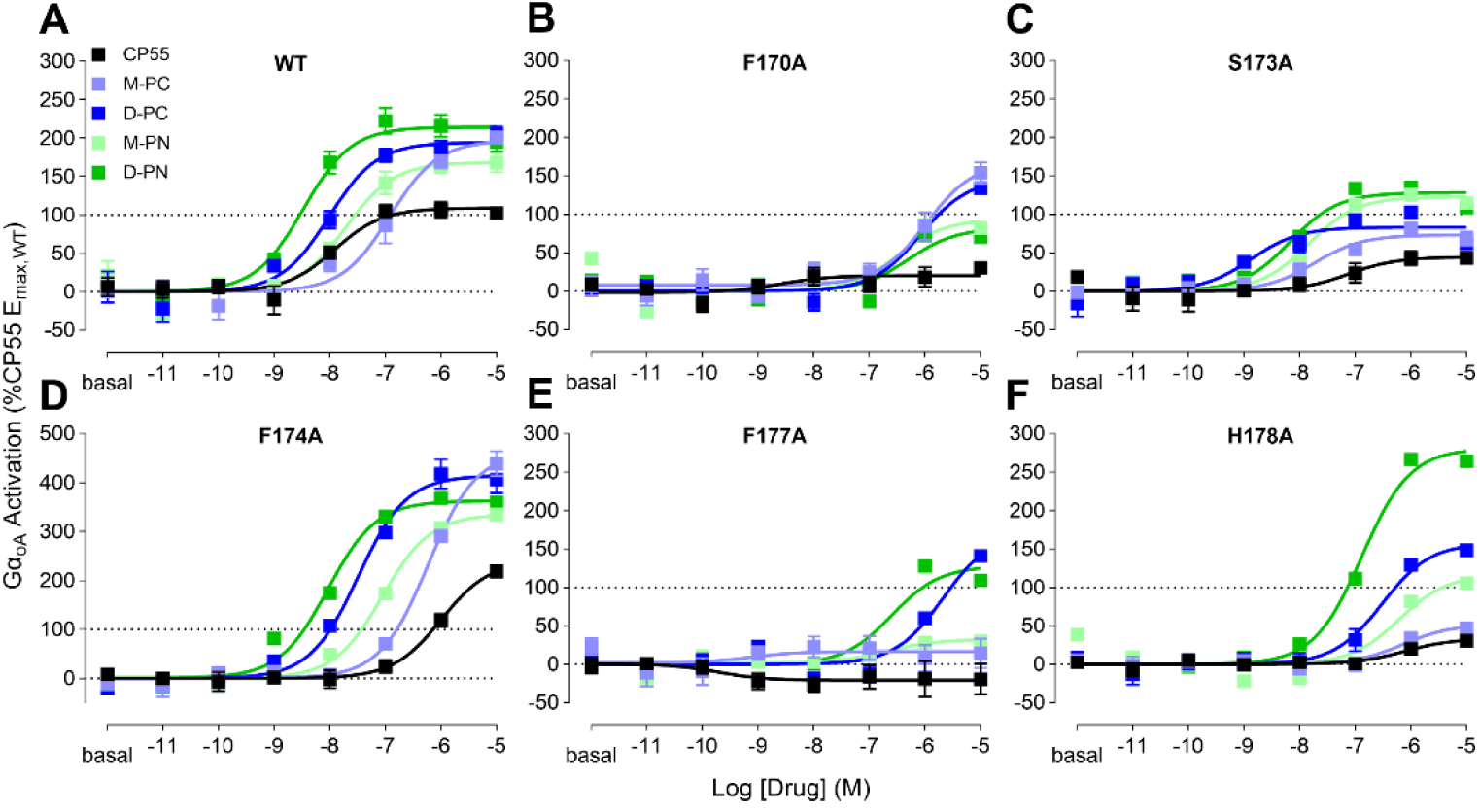
CB1R-TM2 mutational analysis of G-protein activation. Through BRET Gα_oA_ Activation mode, M-PC, D-PC, M-PN, and D-PN are compared to CP55 in the presence of CB1R with the following TM2 mutations: Wildtype (A), F170A (B), S173A (C), F174A (D), F177A (E), H178A (F). Concentration-response curves of SCRA-induced BRET plotted as a percentage of CP55 E_max,WT_. Data presented as means ± SEM of n=4 experiments.

**Supplementary Figure 4:**
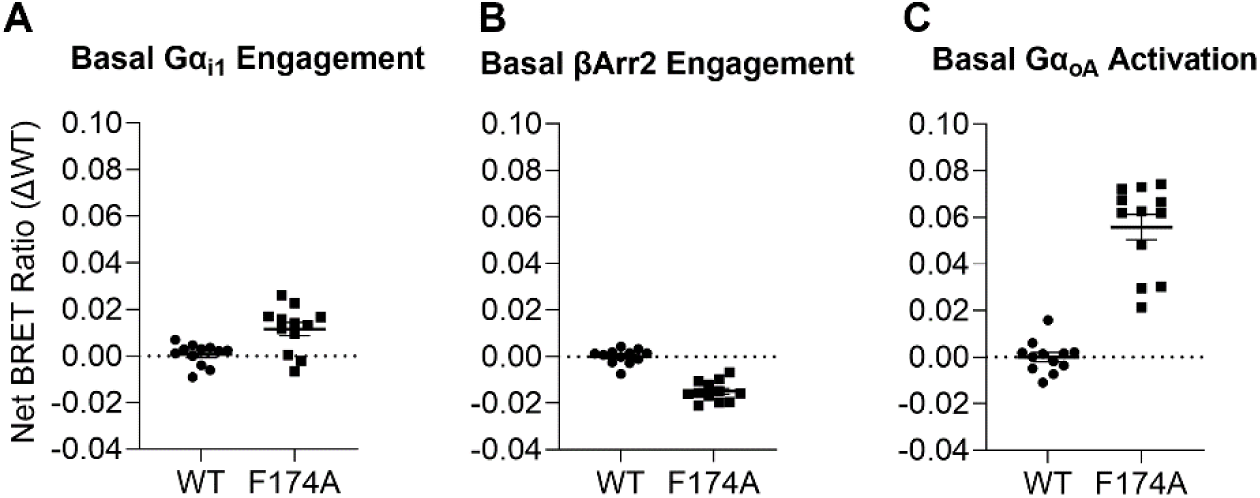
Net BRET Ratios of CB1R F174A in different BRET modes. Calculated basal BRET ratios for CB1R F174A are subtracted from basal BRET ratios of wildtype CB1R for Gα_i1_ Engagement (A), βArr2 Engagement (B), and Gα_oA_ Activation (C) BRET assay modes. Data presented as means ± SEM of n=4 experiments.

**Supplementary Figure 5:**
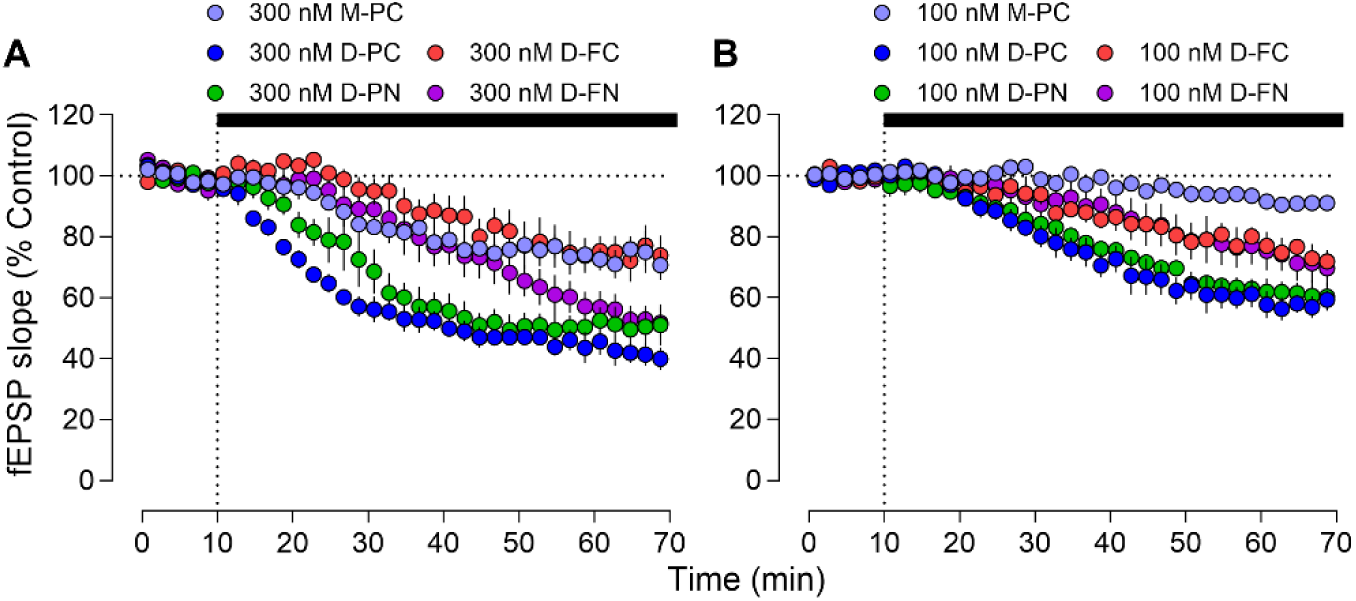
Perfusion of SCRAs at lower concentrations in *ex vivo* slice electrophysiology. Field extracellular postsynaptic potentials (fEPSPs) were recorded every 30 seconds in the CA1 hippocampus during SCRA perfusion. After 10 minutes of baseline, SCRAs were perfused for 60 minutes at 300 nM (A) or 100 nM (B). (N≥4 mice; n≥4 recordings; means ± SEM).

**Supplementary Table 1:** CB1R-TM2 MutationalAnalysis withSCRA Head Moiety (Gα_oA_ Activation)

| Supplementary Table 1: CB1R-TM2 Mutational Analysis with SCRA Head Moiety (G<sub>o</sub>A Activation) |  |  |  |  |  |  |  |  |  |  |  |  |
| --- | --- | --- | --- | --- | --- | --- | --- | --- | --- | --- | --- | --- |
|  | WT |  | F170A |  | S173A |  | F174A |  | F177A |  | H178A |  |
|  | E<sub>max</sub> (% CP55,WT) | pEC<sub>50</sub> (M) | E<sub>max</sub> (% CP55,WT) | pEC<sub>50</sub> (M) | E<sub>max</sub> (% CP55,WT) | pEC<sub>50</sub> (M) | E<sub>max</sub> (% CP55,WT) | pEC<sub>50</sub> (M) | E<sub>max</sub> (% CP55,WT) | pEC<sub>50</sub> (M) | E<sub>max</sub> (% CP55,WT) | pEC<sub>50</sub> (M) |
| CP55 | 100.0 ± 7.03 | 7.92 ± 0.19 | 30.3 ± 6.76 | N.D. | 44.2 ± 8.79* | 7.13 ± 0.49 | 240.1 ± 17.46* | 5.99 ± 0.12* | -19.5 ± 19.90 | N.D. | 33.4 ± 7.10* | 6.12 ± 0.38 |
| M-PC | 197.3 ± 6.43* | 7.91 ± 0.11 | 168.7 ± 18.23* | 5.97 ± 0.19* | 73.1 ± 5.06 | 7.73 ± 0.20 | 463.9 ± 17.87* | 6.23 ± 0.07* | 13.9 ± 18.99 | N.D. | 51.8 ± 8.80 | 6.14 ± 0.31* |
| D-PC | 194.6 ± 8.38* | 8.02 ± 0.14 | 146.4 ± 10.13 | 6.08 ± 0.12* | 83.0 ± 5.14 | 8.88 ± 0.22 | 414.5 ± 11.50* | 7.49 ± 0.08 | 171.7 ± 15.06* | 5.67 ± 0.13* | 157.8 ± 10.35* | 6.51 ± 0.15 |
| M-PN | 168.5 ± 8.18* | 7.71 ± 0.14 | 93.5 ± 21.15 | 6.49 ± 0.50 | 122.7 ± 9.87 | 7.84 ± 0.23 | 335.1 ± 9.12* | 7.05 ± 0.07 | 30.4 ± 11.10 | N.D. | 116.3 ± 15.57 | 6.22 ± 0.26* |
| D-PN | 214.6 ± 7.70* | 8.48 ± 0.12 | 82.1 ± 14.29 | 6.3 ± 0.35* | 128.0 ± 7.42 | 8.17 ± 0.18 | 362.5 ± 9.54* | 8.04 ± 0.08 | 127.8 ± 14.87 | 6.6 ± 0.26 | 280.8 ± 9.83* | 6.88 ± 0.08 |

Data analyzed by one-way ANOVA followed by post-hoc Dunnett test. Data presented as means ± SEM. Significance of \* represents p<0.05 compared to CP55 E<sub>max</sub>, pEC<sub>50</sub> of the wildtype construct. For concentration-response curves with a low potency shift and/or low E<sub>max</sub>, the pEC<sub>50</sub> was not determined (N.D.) and the E<sub>max</sub> shown is the effect at the maximum concentration tested (10 μM).

